# Parallel emergence of perisomatic inhibition and ripples in the developing hippocampal circuit

**DOI:** 10.1101/2025.06.17.660075

**Authors:** Arnaldo Ferreira Gomes Da Silva, Olivier Dubanet, Hervé Rouault, Xavier Leinekugel

**Affiliations:** INMED, INSERM, Aix-Marseille University, Marseille, France; Department of Neuroscience, Kavli Institute for Neuroscience, Wu Tsai Institute, Yale University, New Haven, CT 06510, USA; Aix-Marseille Université, Université de Toulon, CNRS, CPT UMR 7332, Turing Centre for Living systems, Marseille, France

**Keywords:** ripples, hippocampus, perisomatic inhibition, development

## Abstract

During hippocampal Sharp Wave Ripples, sequences of awake coding activities are replayed with a rhythmic timing and high level of synchrony favorable for synaptic plasticity and transfer of information to downstream structures. Previous work reported the emergence of ripples at P10 in the CA1 region, together with the development of inhibition. On the other hand, neither the relationships between perisomatic inhibition and ripples nor their developmental emergence have been investigated in CA3, in which ripples have a different frequency profile (90-110Hz instead of 140-200Hz in CA1), functional perisomatic inhibitory circuits have different properties, and some developmental aspects such as neurogenesis or interneuron maturation are early compared to CA1. We have here investigated the hypothesis of a conjoint and earlier appearance and maturation of ripples and perisomatic inhibition in the CA3 hippocampal region compared to CA1.

We report a parallel sequence of events in CA3 and CA1, starting with the early expression of perisomatic GABAergic synaptic activity combined with the emergence of ripple activity. Interestingly, perisomatic inhibition and ripple activity follow parallel maturation trajectories, beginning in CA3 at P7 with immature (i.e. not functional yet) inhibition and immature ripples (proto-ripples) with labile oscillatory frequency. Mature functional perisomatic inhibition and clear high-frequency ripple activity progressively emerge between P10 and P12, reaching adult-like properties by P13. A similarly progressive maturation of ripples occurs in CA1, from P11 to P15. The progressive emergence of functional inhibition and specific patterns of neuronal activity likely support the progressive emergence of cognitive function to which they are necessary prerequisite.

## INTRODUCTION

Sharp wave ripples (SPW-Rs) are considered essential for learning and memory ^1-3^. During SPW-Rs, sequences of awake coding activities are replayed at fast or real-time scale ^4-9^, in a rhythmic timing that might promote synaptic plasticity and a high level of synchrony that may facilitate summation and transfer of information to downstream structures ^3,10^. Ripples are expressed in both CA3 and CA1, but were studied almost exclusively in CA1, even though CA3 holds a strategic position in the hippocampal circuit, putatively performing pattern completion and being a main source of CA1 input ^3,11-14^.

In CA1, ripples are believed to promote the activation of subicular cells and mPFC as a process of memory transfer ^3^. After long debates regarding their mechanism of generation ^15-17^, recent evidence suggests that they are generated by the local interplay between interneurons and glutamatergic pyramidal cells. PV basket cells would provide perisomatic IPSPs with a frequency and time course responsible for the 150-200Hz oscillation expressed by their target pyramidal cells ^18,19^. Previous work also suggests that perisomatic inhibition and ripples would co-emerge in CA1 during development. Dard et al. reported a surge in perisomatic GABAergic innervation and the appearance at P9-10 of functional inhibition of pyramidal cells locked to the occurence of animal body movements ^20^. Earlier studies reported the emergence of ripples at P10 in the CA1 region ^21^, which were recently shown to correlate with the development of inhibition and to be prevented by the chemogenetic blockade of GABAergic interneurons ^22^. On the other hand, neither the relationships between perisomatic inhibition and ripples or their developmental emergence have been investigated in CA3, in which ripples have a different frequency profile (90-110Hz instead of 140-200Hz in CA1) ^23,24^, functional perisomatic inhibitory circuits have different properties ^25,26^, and some developmental aspects such as neurogenesis ^27,28^ or interneuron maturation ^29-32^ are early compared to CA1. We have here investigated the hypothesis of a conjoint and earlier appearance and maturation of ripples and perisomatic inhibition in the CA3 hippocampal region, compared to CA1.

### Progressive emergence of time-locked perisomatic inhibition in CA3

In order to investigate the developmental emergence of perisomatic inhibition in CA3 during spontaneous activity in vivo, we have taken advantage of the expression of fIPSPs as a readout of PV-mediated inhibitory GABAergic synaptic events. fIPSPs were not observed until P7 in the CA3 hippocampal region of urethane-anesthetized mice. As illustrated in Figure 1B and C, no spontaneous fIPSPs were detected before P7 (n=5 mice at age P5-6). The first detectable fIPSP events appeared at P7, and their frequency of occurrence increased with age from about 30 events/min at P7 (median 30.5 events/min, IQR 24.1-32.6, n=3 mice) to more than 670 events/min in adult (median 676.8 events/min, IQR 628.1-793.8, n=6 mice). As shown in Figure 1E, analysis of cross-correlograms between fIPSPs and multi-unit activity (MUA) revealed 3 distinct developmental phases. First, between P7 and P9 (n=7 mice), MUA did not appear significantly affected by fIPSPs. P10 to P12 appeared as a transition period, during which half of the recordings displayed inhibited activity following fIPSPs (i.e. firing rate below [baseline - 2SD] from 0 to at least 4ms after fIPSP-peak, cf STAR METHODS) while the other half was globally not affected (n=10 mice). The reasons for a progressive maturation of the inhibitory action of fIPSPs are unclear. It may be that some individual cells are inhibited and others excited, resulting in a global apparent absence of effect. On the other hand, all recordings showed adult-like time-locked inhibition from P13 onwards (n=10 P13-18 and 6 adult mice). These results suggest that perisomatic GABAergic transmission emerges gradually in the CA3 region between P7 and P13 when it reaches mature functional inhibitory action over the CA3 circuit. We hypothesize that the emergence of fIPSPs at P7 in CA3 corresponds to a surge in perisomatic GABAergic innervation, as described in CA1 a couple of days later (P9) ^20^.

**Figure 1.**
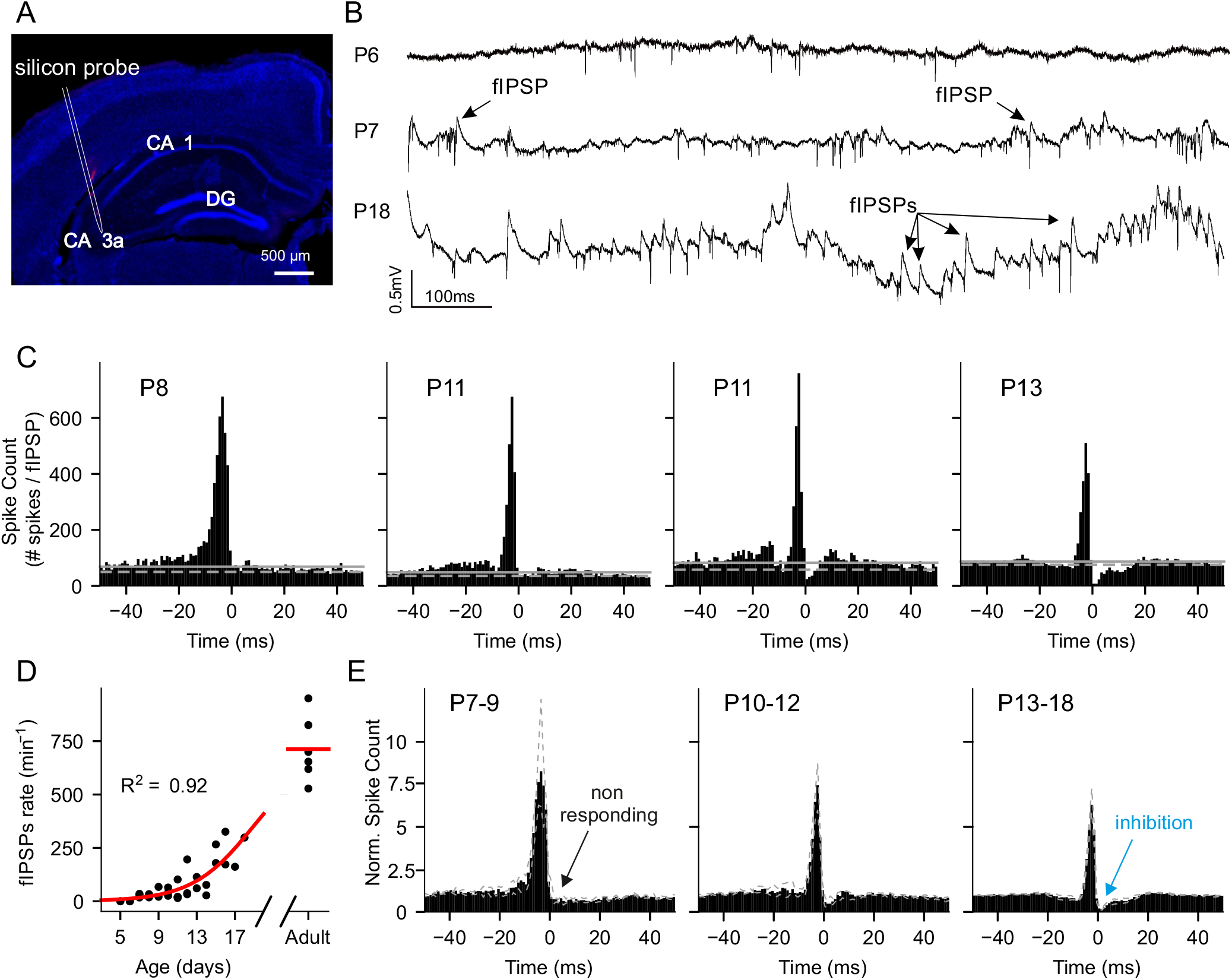
Developmental emergence of fIPSPs in CA3. A. Post-hoc histological verification of recording location showing the electrode track in the CA3a hippocampal region of a vertical brain section (red, [DiI]-labeled silicon probe; blue, DAPI staining). B. Example traces of spontaneous neuronal activity (wide-band field-recording, 0.1Hz–9KHz) in the CA3a pyramidal layer at ages P6, P7 and P18. C. Example peri-event time histograms between multi-unit activity and fIPSPs (reference) from individual animals at ages P8, P11 and P13 (plain horizontal line, mean baseline firing, dotted line mean -2SD). Note the emergence of functional perisomatic inhibition (post-fIPSP inhibitory trough) at P11 (middle right histogram), generalized to all animals from P13 onwards. D. fIPSP rates of occurrence throughout development. E. Average normalized peri-event time histograms (mean, plain bars, dotted lines, +/- SD, bin 1ms) between multi-unit activity and fIPSPs (reference) at ages P7-9 (n=7 mice), P10-12 (n=10 mice) and P13-18 (n=10 mice), illustrating the lack of neuronal response at P7-9 and clear inhibition from P13 onwards.

### Parallel emergence of ripples and perisomatic inhibition during the second postnatal week in CA3

Since in CA1 the emergence of ripples was associated with the developmental expression of perisomatic inhibition, which we observed to occur at P7 in CA3, we hypothesized that ripples may also emerge at P7 in CA3. Typically, CA3 ripples are identified as collective bursts of activity associated with a prominent field potential oscillation between 80 and 110 Hz. As illustrated in Figure 2, we indeed observed transient high frequency oscillations in the ripples range from P7 onwards (median 0.14 events/min, IQR 0.09-0.55, P7-9: n=8 mice) but not before, which frequency of occurrence increased with age. On closer examination however, it turned out these events had a rather heterogeneous time-frequency profile. While some presented the typical high frequency power band well separated from background activity, others were less clearly defined or confined in terms of frequency domain, even though clearly oscillatory. The related spike discharge appeared also less clearly time-locked to the field oscillations. Interestingly, the expression of these events appeared strongly developmentally regulated, and while the dominant class of events at P7-9 (median 82.8 %, IQR 80-85.7, n=8 mice), they progressively disappeared between P10 and P12 (n = 13 mice), so that from P13 onwards almost all events had a classical adult-like ripple time-frequency profile (P13-18: median 2.6 %, IQR 0.6-7.5, n=12 mice; P7–19 vs P13–18: p<0.001).

**Figure 2.**
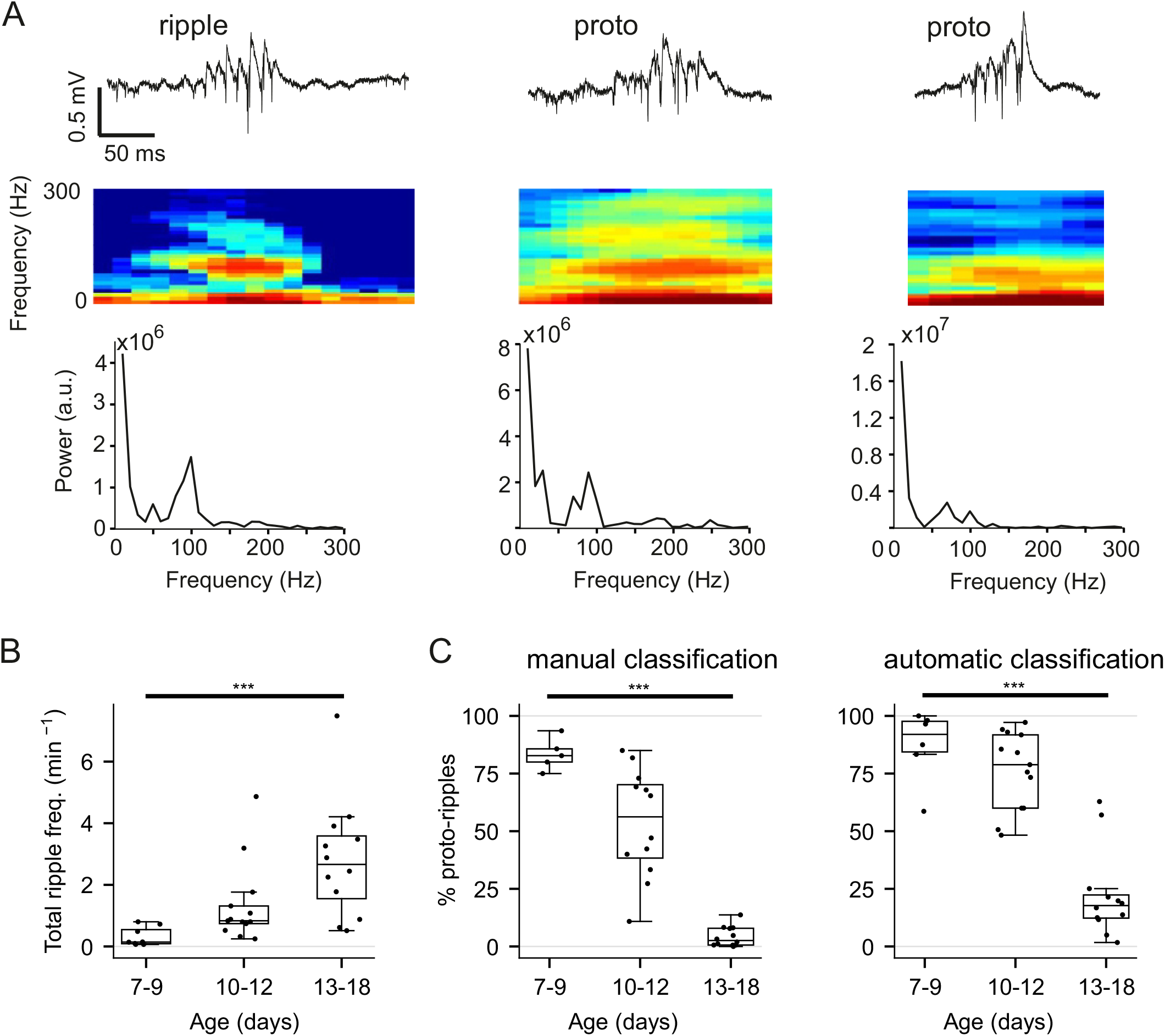
Developmental emergence of ripples in CA3. A. Example traces (top, wide-band LFP, 0.1 Hz-9 KHz), time frequency spectrogram (middle) and power spectral density (bottom) of an individual ripple (left) and 2 individual proto-ripples (middle and right) recorded in the CA3a pyramidal layer at developmental stage P7-12. B. Frequency of occurence of all detected ripples (including proto-ripples) across ages (P5-6, n=5 mice, P7-9, n=8 mice, P10-12, n=13mice, P13-18, n=12 mice). C. Proportion of proto ripples (relative to all detected ripples, including proto ripples) determined by manual (left) or automatic ripple and proto-ripple sorting. ***p<0.001, box plots, median and quartiles.

In order to have an objective distinction and quantification of these two kinds of events, that we distinguished as proto-ripples vs classical ripples, we trained an artificial neural network to discriminate and identify ripples and proto-ripples. The network demonstrated robust classification performance, as evidenced by the confusion matrix (Supplementary Fig. 1C) with an overall accuracy of 83%, a proto-ripple sensitivity of 70% and a ripple sensitivity of 78%. A UMAP analysis of the network’s 4-dimensional output vectors into a 2D space (Supplementary Fig. 1B) revealed an interesting organization. While the overall UMAP formed a single connected component, the proto-ripple cluster was not positioned intermediately between the ripple and background clusters. Instead, the axis defining ripple/proto-ripple clustering appeared orthogonal to the axis separating ripple/proto-ripple from background. This arrangement suggests that proto-ripples are not merely noisy signals, but rather an emergent feature distinct from general background activity. Furthermore, by coloring the UMAP plots for proto-ripples and ripples according to animal age (Supplementary Figure 1D), we observed a clear progression, with a continuous gradient between the proto-ripple and ripple clusters. This spatial arrangement within the UMAP, coupled with the age-dependent gradient, strongly supports the hypothesis that proto-ripples represent an earlier developmental stage in the emergence of mature ripples. Moreover, following the frequency of ripples and the ratio proto-ripple vs. ripples (Figure 2B and Supplementary Figure D), it appears that ripple-like activity closely matches the emergence and maturation of perisomatic inhibition in CA3, with first expression at P7 as immature (proto-ripples and fIPSPs without clear functional inhibition of the circuit), then a transition period from P10 to P12 with a mixture of immature and mature events, and finally a mature state from P13 onwards.

### Delayed and progressive emergence of ripples in CA1

Since CA3 has been proposed in former studies to have an earlier maturation than CA1 but along the same path, we hypothesized that the CA1 circuit as well may show progressive maturation of ripple activity. As in earlier studies ^21,22^, we did not observe any ripple activity in CA1 until P10 (n=4 mice at P9-10, cf. Figure 3). But as in CA3, we observed the developmentally regulated expression of proto-ripples and classical ripples, between P11 and P15 (cf Supplementary Figure 1E and Figure 3C). At P11-12 (n=7 mice), nearly all ripple events displayed immature features (proto ripples) in terms of time frequency signature (median 91.18%, IQR 81-100% of events were proto ripples, n=7 mice), while after a transition period between P13 and P14 (n=4 mice) with a mixture of proto and classical ripples (median 31.65%, IQR 22.2-46.8% of proto ripples, n=4 mice), most ripple events appeared with a classical adult-like signature, from P15 onwards (median 8.4%, IQR 6.6-16.6% of proto ripples, n=8 mice; P11-12 vs P15-P18: p<0.01).

**Figure 3.**
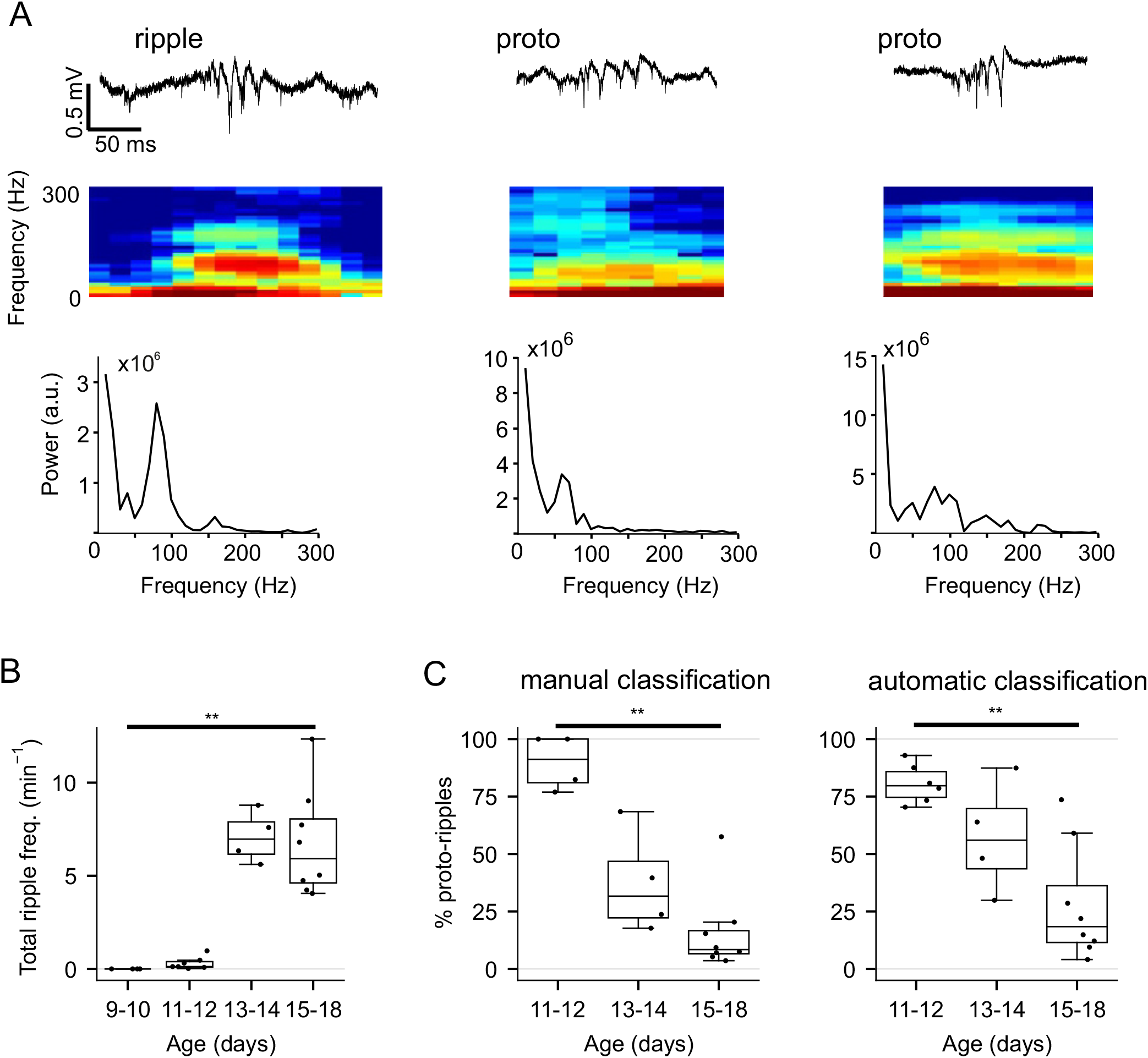
Developmental emergence of ripples in CA1. A. Example traces (top,wide-band LFP, 0.1 Hz-9 KHz), time frequency spectrogram (middle) and power spectral density (bottom) of an individual ripple (left) and 2 individual proto-ripples (middle and right) recorded in the CA1 pyramidal layer at P11-13. B. Frequency of occurence of all detected ripples (including proto-ripples) across ages (P9-10, n=4 mice, P11-12, n=7 mice, P13-14, n=4mice, P15-18, n=8 mice). C. Proportion of proto ripples (relative to all detected ripples, including proto ripples) determined by manual (left) or automatic ripple and proto-ripple sorting. **p<0.01, box plots, median and quartiles.

### Conclusion

We therefore report that in CA3, as in CA1, perisomatic inhibition emerges concomitantly with significant functional changes such as the expression of ripple activity. Although the CA3 circuit appeared to undergo earlier maturation than that of CA1, as suggested by previous studies, we now report a parallel sequence of events starting with the early expression of perisomatic GABAergic synaptic activity combined with the emergence of ripple activity. Interestingly, both perisomatic inhibition and ripple activity follow parallel maturation trajectories, beginning in CA3 at P7 with a lack of apparent inhibition in the circuit, while proto-ripples exhibit labile frequency expression. A transition period occurs between P10 and P12, during which mature functional perisomatic inhibition and ripple activity progressively emerge, both reaching adult-like properties by P13. Further experiments directly manipulating specific populations of GABAergic interneurons are awaited to evaluate in which respect CA3 ripples directly depend on perisomatic inhibition and more specifically PV interneurons, but the concomitant emergence and maturation of fIPSPs and ripples during development suggests that it may be the case in CA3 as it was demonstrated for the CA1 circuit.

## ACKNOWLEDGEMENTS

This work has received support from the French government under the “France 2030” program via A*Midex (Initiative d’Excellence d’Aix-Marseille Université, AMX-19-IET-004, ANR-17-EURE-0029, ANR-16-CONV-0001), INSERM (XL), Région Nouvelle Aquitaine (XL), Agence Nationale pour la Recherche (XL: ANR-18-CE16-0015-02, ANR-18-CE16-0020-04) and Institut pour la Recherche en Santé Publique (XL: IReSP AAP-2023-EIMA-332751). We thank all the personnel of the animal facilities of the NeuroCentre Magendie and of INMED for animal care.

## AUTHOR CONTRIBUTIONS

XL initiated and designed the study. OD and AFGDS performed the experiments. OD, AFGDS, HR and XL analyzed the data. AFGDS, HR and XL prepared the figures and wrote the manuscript. All authors have read and edited the manuscript.

## STAR METHODS

### EXPERIMENTAL MODEL DETAILS

#### Animals

In this study, we used a total of 45 (33 males and 12 females) C57BL/6 PV-Cre transgenic mice (Jackson Laboratory, B6;129P2-Pvalbtm1(cre)Arbr/J)^33^, including juveniles (5 to 18 postnatal days, P5-18) and adults (8-10 weeks old). Animals were kept on a 12h:12h light/dark cycle and provided with ad libitum access to food and water. All experimental procedures were conducted in accordance with the EU directives regarding the protection of animals used for experimental and scientific purposes (86/609/EEC and 2010/63/EU) and approved by the French Ministry for Research, after ethical evaluation by the local Ethics Committee Bordeaux - CEEA50.

### METHOD DETAILS

#### Surgical procedure and multi-site extracellular recording *in vivo*

Extracellular recordings of spontaneous multi-unit activity and local field potentials were performed from the right dorsal hippocampal CA1 and CA3a areas of juvenile (P5-18) and adult (8-10 weeks) mice anesthetized with urethane (Sigma-Aldrich) and a mixture of ketamine (Imalgene 1000, Merial) and xylazine (Rompun 2%, Bayer). After a subcutaneous injection of Rimadyl (5 mg/kg for adult, 1 mg/kg for juvenile mice) 30 minutes before the surgery to prevent brain inflammation, the mice were anesthetized with urethane, according to age: 0.75g/kg IP complemented with half the initial dose whenever necessary (typically every 30 min) for juvenile animals weighting less than 8 grams; 1.7g/kg IP complemented with ketamine/xylazine (1.3mg/kg and 0.13mg/kg, IM, respectively) as needed, typically every 30 minutes for juvenile animals weighting more than 8 grams; 1.7g/kg urethane IP supplemented with ketamine/xylazine (IM, respectively 6.6mg/kg and 0.66mg/kg) as needed, typically every 30 to 60 minutes for adult animals. The mouse was then placed in a stereotaxic frame (David Kopf Instruments), kept on a thermal blanket (Physitemp) to maintain body temperature at 37.5°C. The eyes were covered with a protective liquid gel (Ocrygel). The scalp and periosteum over the dorsal surface of the skull were removed. Reference and ground wires were implanted in the cerebellum and attached to a 2-pin connector, maintained together with a small custom-made horizontal stainless-steel bar anchored to the skull above the cerebellum with dental acrylic. A thin anchoring layer of dental acrylic (Super-Bond, Frapident) was applied on the exposed skull, except above the right CA3 and CA1 regions of the hippocampus for silicon probe insertion and recording. After performing the craniotomy at coordinates -1.2 to -1.6mm AP, +1.5 to +2.5mm ML, the dura was gently removed and a multi-site silicon probe (Neuronexus Technologies, Buzsaki-16, 2 shanks separated by 200 μm, each with 8 recording sites, 20 μm separation), covered with DiI (Molecular Probes) for post-hoc verification of electrode position, was inserted with an angle of 15° from vertical through the neocortex until the pyramidal layer of the dorsal CA3a hippocampal region (depth of 1.2 to 1.7 mm). The stratum pyramidale was recognized by the presence of multiple firing cells and fIPSPs, and confirmed with the post-hoc identification of DiI labeling.

For CA1 recordings, a distinct silicon probe (Neuronexus Technologies, BuzLin32, 3 shanks separated by 200 μm, with 8 recording sites each separated by 20 μm on the two lateral shanks and 16 channels on the middle shank arranged vertically with a 50 μm vertical separation), also labeled with DiI, was inserted vertically through the neocortex to the pyramidal layer of the dorsal CA1 region, at coordinates 1.2 to -1.6mm AP, +1.2 to +1.6mm ML. The stratum pyramidale (depth 1.0 to 1.5 mm) was recognized by the presence of multiple firing cells and SPW-ripples, and confirmed with the post-hoc identification of DiI labeling.

Recordings were performed using an integrated Digital Lynx SX system (Neuralynx, Cheetah v5.0 software, 24bits, sampling rate 32KHz, bandpass 0.1Hz to 9KHz, range ± 5mV for LFP analysis, as well as bandpass 1Hz to 9KHz, range ± 1mV for spike sorting), and stored on a PC for offline analysis. The recording started approximately 15 minutes after the insertion of the probe.

The spikes were detected and extracted with SpikeDetekt, automatically clustered with KlustaKwik2, and the resulting clusters were manually verified and refined with KlustaViewa using the KlustaSuite software (Rossant et al., 2016) (https://www.ucl.ac.uk/cortexlab/). Because the number of identified units was too small to draw statistically relevant conclusions, we decided to analyse only the multi-unit activity (MUA).

#### Histology

After recording, the brain was removed and incubated in paraformaldehyde (PFA) 4% during 24h for post-fixation. Coronal sections (70μm-thick, cut using a VT1000S Leica vibratome) were prepared to verify the position of the DiI-coated electrodes, mounted and coverslipped with DAPI Fluromount-G. Epifluorescence images were obtained with a Zeiss AxioImager Z2 microscope.

### QUANTIFICATION AND STATISTICAL ANALYSIS

#### Detection and effects of perisomatic GABAergic events

The fIPSPs were detected as previously described^25^, with the MiniAnalysis software (6.0.3 Synaptosoft Inc.) from the raw signal using the following parameters (manually adjusted within the indicated ranges []): amplitude (mV), min = [150, 450], rise time (ms), max = [2.8, 5], area, min = [0,1000], max = [5000, max], halfwidth (ms) min = [0.9, 2], max = [10, max]. Peri-event time histograms display the time-distributions of neuronal activity relative to fIPSPs detected from the same electrode/shank. The number of spikes was counted within time-bins of 1ms around fIPSP-peak taken as reference events, and normalized by the number of events, resulting in spike count per fIPSP event (cf Figure 1C). Baseline activity was computed as the mean (and SD) of values between -50 and -25ms. Neuronal activity was considered as inhibited by fIPSP-associated events if the firing rate (MUA) decreased and remained below that of [baseline - 2SD] from 0 to at least 4ms after fIPSP-peak. Neuronal activity was considered as excited by fIPSP-associated events if the firing rate increased above [baseline + 3SD] within 4ms after fIPSP-peak. Neuronal activity was considered as non-responding if the firing rate (MUA) was neither inhibited nor excited by fIPSP-associated events.

#### Ripples and proto-ripples detection, quantification and statistical analysis

Data were visualized and processed using NeuroScope and NDManager from the Neurosuite software (Hazan et al., 2006) (http://neurosuite.sourceforge.net), and analyzed using Origin (OriginLab, Northampton, MA) and MATLAB (MathWorks) built-in or custom-built procedures.

Ripple / proto-ripple detection was performed by band-pass filtering (80-150 Hz range for CA3, 80-200 Hz for CA1), squaring and normalizing, then thresholding the field potential recorded in the pyramidal layer. Ripples / proto-ripples were defined as events peaking at >7 standard deviations and lasting <500 ms. Automatically detected events were then manually classified into three categories: ripples, proto-ripples and false positives (background). Comparisons between groups were evaluated using the Kruskal-Wallis and Mann Whitney tests. Differences were considered statistically significant at p<0.05.

#### Neural Network Training

A deep neural network (see supp. fig. 1) was implemented using PyTorch and trained with the Lightning framework to differentiate between four categories of 128ms LFP segments: automatically detected events manually annotated as either “ripple”, “proto-ripple” or “background”, as well as a set of randomly selected segments from non-annotated sections of the recording (“random”).

To build the dataset, a 200ms window centered on the peak power of automatically detected (and manually annotated as ripple / proto-ripple / background) events was extracted, from which a 128 ms window was extracted and fed into the deep neural network. The extracted LFP segments (full dataset: n=5051 “ripple” segments, 1317 “proto-ripple” segments, 18712 “background” segments and 25080 “random” segments), were divided into training (80%) and validation (20%) sets. To address class imbalance, a weighted sampling strategy was employed during training, ensuring that an equal number of samples were drawn from each class.

The network was trained for 600 epochs using a cross-entropy loss function. After training, the network’s performance was evaluated on the validation set to assess its ability to accurately classify LFP segments into background, proto-ripple, and adult ripple categories based on the manual annotations. The “random” category was used only for the training, and subsequently ignored.

Source code:

For the cross-correlogram analysis and supervised ripple classification, scripts were written in Python and available at https://gitlab.com/rouault-team-public/ferreira_et_al.

## SUPPLEMENTARY INFORMATION

**Supplementary Figure, related to: Artificial neural network based classification of hippocampal ripples and proto-ripples**.

A. Schematic of the convolutional/dense neural network architecture employed for the classification of local field potential (LFP) segments into background activity, proto-ripples, or ripples. Numbers of channels and sizes of the 1D signals are indicated.

B. Two-dimensional UMAP embedding of the 4-dimensional feature space extracted by the neural network for all analyzed LFP input signals.

C. Confusion matrix evaluating the accuracy of the supervised classification model on the held out validation dataset.

D. Developmental trajectories of proto-ripples and ripples in CA3. Left, UMAP representation: green circles, events manually labeled as ripples; red circles, events manually labeled as proto-ripples; inside color according to age (postnatal day, cf. color bar). Right, total ripple frequency of occurrence and proportion of proto-ripples in the validation set (20% of events not included in the artificial network training, which explain the total frequency about 5 times lower than the total frequencies in the whole dataset presented on main figures 2 and 3.).

E. Developmental trajectories of proto-ripples and ripples in CA1. Same as in D but for CA1 events.

