## Supplemental-Figure for "Parallel emergence of perisomatic inhibition and ripples in the developing hippocampal circuit"

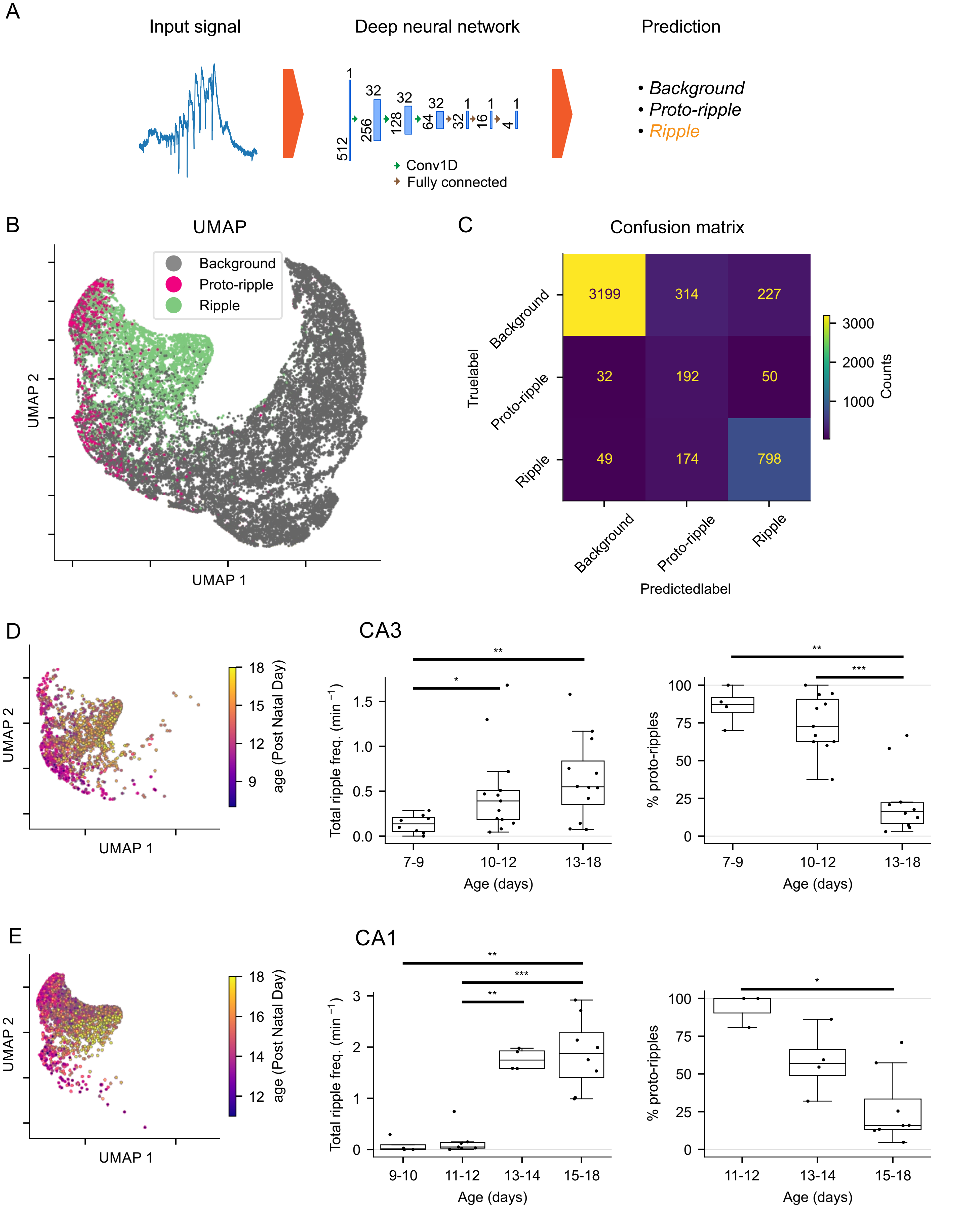


**Supplementary Figure, related to: Artificial neural network based classification of hippocampal ripples and proto-ripples.**
A. Schematic of the convolutional/dense neural network architecture employed for the classification of local field potential (LFP) segments into background activity, proto-ripples, or ripples. Numbers of channels and sizes of the 1D signals are indicated.
B. Two-dimensional UMAP embedding of the 4-dimensional feature space extracted by the neural network for all analyzed LFP input signals.
C. Confusion matrix evaluating the accuracy of the supervised classification model on the held out validation dataset.
D. Developmental trajectories of proto-ripples and ripples in CA3. Left, UMAP representation: green circles, events manually labeled as ripples; red circles, events manually labeled as proto-ripples; inside color according to age (post natal day, cf. color bar). Right, total ripple frequency of occurrence and proportion of proto-ripples in the validation set (20% of events not included in the artificial network training, which explain the total frequency about 5 times lower than the total frequencies in the whole dataset presented on main figures 2 and 3.).
E. Developmental trajectories of proto-ripples and ripples in CA1. Same as in D but for CA1 events.
